# HOW TO MEASURE THE INFLUENCE OF LANDSCAPE ON POPULATION GENETIC STRUCTURE: DEVELOPING RESISTANCE SURFACES USING A PATTERN-ORIENTED MODELING APPROACH

**DOI:** 10.1101/2020.02.20.958637

**Authors:** Kelly Souza, Jesús N. Pinto-Ledezma, Mariana Pires de Campos Telles, Thannya Nascimento Soares, Lazaro José Chaves, Clarissa Bonafé Gaspar Ruas, Ricardo Dobrovolski, José Alexandre Felizola Diniz-Filho

**Affiliations:** Programa de Pós-Graduação em Genética, PGBM, ICB, Universidade Federal de Goiás. Campus II (Samambaia), Goiânia, GO, Brasil; Department of Ecology, Evolution and Behavior, University of Minnesota, 1479 Gortner Ave, Saint Paul, MN 55108, USA; Departamento de Genética, Universidade Federal de Goiás. Campus II (Samambaia), Goiânia, GO, Brasil and Escola de Ciências Agrárias e Biologia, PUC-GO, Av. Engler, 286-316 - Jardim Mariliza, Goiânia, GO, Brasil; Departamento de Genética, Universidade Federal de Goiás. Campus II (Samambaia), Goiânia, GO, Brasil; Escola de Agronomia, Universidade Federal de Goiás. Campus II (Samambaia), Goiânia, GO, Brasil; Graduanda em Ciências da Computação, Universidade Estadual Paulista, Rio Claro, SP, Brasil; Departamento de Zoologia, Universidade Federal da Bahia, Salvador, BA, Brasil; Departamento de Ecologia ICB, Universidade Federal de Goiás, Goiânia, GO, Brasil

**Keywords:** gene flow, FST, *Dipteryx alata*, *c*ircuitscape, mantel

## Abstract

There are several approaches to understand how a landscape, with its several components, affects the genetic population structure by imposing resistance to gene flow. Here we propose the creation of resistance surfaces using a Pattern-Oriented Modeling approach to explain genetic differentiation, estimated by pairwise FST, among “Baruzeiro” populations (*Dipteryx alata*), a tree species widely distributed in Brazilian Cerrado. To establish the resistance surface, we used land use layers from the area in which the 25 “Baruzeiro” populations were sampled, generating 10000 resistance surfaces. To establish the resistance surface, we used land use layers from the area in which the 25 “Baru” populations were sampled, generating 10000 resistance surfaces. We randomized the cost values for each landscape component between 0 and 100. We use these surfaces to calculate pairwise matrices of the effective resistance among populations. Mantel test revealed a correlation of pairwise FST with a geographical distance equal to r = 0.48 (P < 0.001), whereas the Mantel correlations between pairwise FST and the generated resistance matrices ranged between r = −0.2019 and r= 0.6736. Partial regression on distance matrices was used to select the resistance matrix that provided the highest correlation with pairwise FST, based on the AIC criterion. The selected models suggest that the areas with lower resistance are characterized as natural savanna habitats of different forms, mainly arboreal dense savannas. In contrast, roads, big rivers, and agricultural lands cause higher resistance to gene flow.

## Introduction

Landscape structure can influence ecological processes, such as dispersal, migration and flow gene, at different levels of biological organization and spatial scales, including those driving the population genetic structure (Manel et al. 2003; Koffi et al. 2007). For example, a strong genetic structure (i.e., high differentiation among demes or local populations) appears when habitat loss decreased connectivity and, therefore, the dispersal capacity, dividing populations and disrupting gene flow (King and With 2002, Amos et al., 2012; Braga et al. 2019). This decrease in gene flow results in loss of genetic variation and inbreeding depression, most likely increase the probability of local extinctions (Storfer 1999), changing aspects of species’ life history (Kramer et al. 2008) and eventually reducing its evolutionary potential (Frankham et al. 2004). The analysis of all these processes is the primary goal of a field that quickly developed and advanced in the last 15 years called “Landscape Genetics” (Manel et al. 2003; Holderegger and Wagner 2006; Storfer et al. 2007; Manel and Holderegger 2013). Landscape genetics seeks to evaluate the interaction between landscape features and microevolutionary processes such as gene flow, selection, and genetic drift, integrating, thus geographical, ecological, and genetic information (Manel et al. 2003; Storfer et al. 2007).

In landscape genetics, it is possible to check and interpret the effective distance between individuals or populations, taking into account landscape properties that wild better reflect gene flow (Mateo-Sánches et al. 2015). This relation can be calculated through the “Isolation by Environment” (IBE) models, which describes a pattern in which genetic differentiation increases with environmental differences, independent of geographical distances (Sexton et al. 2014, Jenkins et al. 2010, Wang and Bradburd 2014). Another way to check the effective distance in landscape genetics is by Isolation by Resistance – IBR (McRae 2006), where the distance calculation incorporates the degree of “permeability” of the different landscape components (e.g., forests, croplands, roads) to the dispersion of individuals throughout the landscape. Therefore, this permeability is related to how the landscape affects the movement of organisms between the areas with resources, in terms of biological, physiological, and behavioral characteristics, thus controlling the natural flows of species (Metzger and Ddcamps 1997; Tischendorf and Fahrig 2000). IBE and IBR researchers tend to concentrate on describing patterns, without necessarily investigating the mechanisms that have generated these patterns (Wang and Bradburd 2014). However, all these models are generalized versions of the much older (and simpler) Isolation-by-Distance (IBD) proposed by Sewall Wright in the early 1940’s, in which it is possible to predict an exponential decrease of genetic distance as geographic distances increase, by a balance between dispersal and local genetic drift (see Wright 1943). So, both IBR and IBE can be viewed as more complex cases of IBD in terms of dispersal routes and changes in a balance due to landscape features.

Resistance surfaces have been used to understand how landscape components influence the connectivity among species populations. (Spear et al. 2010; Koen et al. 2012). These surfaces are representations of the degree of connectivity that is attributed to the original landscape components (i.e., considering the organisms of interest) that are used to model their movement through the landscape (Spear *et al.* 2010; Taylor et al.,1993; Coulon et al., 2004; Vignieri, 2005). A crucial step in the development of these surfaces is to attribute values, or costs, to each of the landscape components. Such parameterization will determine how users will be this resistance to model species movement throughout the landscape (Spear et al. 2010; Koen et al. 2012). However, this attribution of costs is, in many cases, subjective and is not based on strict knowledge of species’ traits.

To describe the resistance that the landscape imposes to the gene flow between populations and to reveal information on the processes behind the population’s genetic structure observed patterns, it is possible to use a Pattern-Oriented Modeling technique (Grimm 1994; Grimm et al. 1996; Grimm et al. 2005; Diniz-Filho et al. 2014). The POM provides a conceptual framework to assess the applicability of models by comparing the patterns generated by the model to observed patterns (Kang e Aldstadt, 2018). One can use computational procedures to create a conceptual framework and find the set of parameters that generates the best models replicating an empirical pattern. The creation of this models resulted in the improvement of the quality of the model and the overall understanding of the system (Kang e Aldstadt, 2018; Diniz-Filho et al. 2014), which allows a biological and ecological interpretation of this best set of parameter values (Wiegand et al. 2003).

The system we are interested in refers to the landscape influence in the genetic structure of *Dipteryx alata* populations in Central Brazil. Assuming that genetic diversity has a positive relationship with the resistance landscape (McRae 2006), we used here genetic diversity to attribute the values of resistance of the different landscape components to build resistance surfaces that better explain the genetic structure pattern among populations. The main goal is to understand which landscape features (or combination of them) better define the genetic divergence between populations. This statistical definition minimizes the arbitrariness in the parameterization of these surfaces, so we used the POM approach in the search for matches between simulated and observed patterns (Diniz-Filho et al. 2014). Several lines of evidence suggest that anthropogenic features affect the connectivity and the gene flow in natural landscapes (Pérez-Espona 2008, Ayran et al. 2017; Okamiya and Kusano 2019). Thus, we expect that anthropogenic features will be selected for the POM and present a high resistance for the gene flow. We also hope that landscape features, especially those of the species natural habitat and their dispersers, show less resistance to the gene flow.

## Materials and Methods

For our analyses, we sampled *Dipteryx alata Vog* in 25 localities (local populations hereafter), a tree species widely distributed in the Brazilian Cerrado, popularly known as “Baru” tree or “Baruzeiro.” We estimated the genetic variation of 644 individuals collected, with sample sizes ranging from 12 to 37 individuals in each local population, covering most of the geographical distribution of the species (Table S1), as detailed described elsewhere (especially Soares et al. 2012; Diniz-Filho et al. 2012; Collevatti et al. 2013). Seven microsatellite loci were used to estimate the genetic divergence between pairs of populations using Wright’s F_ST_, calculated using the *pp.fst* function in the *hierfstat* package (Goudet 2005), with modifications on R platform (R Core Team 2014).

We define the study area with a polygon covering all populations, marginally buffered by 20 kilometers around the local populations, covering an area of approximately 1.25 × 106 km^2^. We used the contemporary habitat configuration to define landscape elements to build the resistance surfaces. The overall reasoning is to determine the influence of a landscape modified by changes in the Cerrado region over the last 50 years due to agricultural expansion and population growth in the region (Klink and Machado 2005; Klink 2013). Initially, we adopted land use layers with land cover classes (hydrography, roads, urban areas, and vegetation types) obtained from the Land Cover Maps Biomes, on a spatial scale of 1:250.000, based on Landsat 7 ETM + with pictures of 2002 and Brazilian roads map (see www.mma.gov.br and www.dnit.gov.br). We reclassified the raster maps with approximately 0.02 decimal degrees of resolution into distinct natural (vegetation types of forests, such as savannas, grasslands, and drainage network) or anthropic classes (agricultural areas, urban areas, and roads), (see Table 2 for a description) and calculated the percentage of each class in the landscape. All this information (landscape components) was gathered into a single layer and used to calculate the resistance surfaces. For our analysis, when a cell has two or more classes, the class with higher occurrence was selected to describe the cell, using ArcGIS 9.3 software (ESRI, 2011).

### Landscape Analysis and Pattern-Oriented Modeling (POM)

The landscape resistance to the gene flow of a species is caused by the interaction between its biological traits for dispersion (or their dispersers) and all land cover classes structured in a resistance surface. The resistance surface is the numerical demonstration of the quantity of resistance imposed on the gene flow among the populations on the study. One of the major challenges in Landscape Genetics is the assignment of resistance values to this landscape components because its interaction (and usually primary biological data) is unknown (Spear et al. 2010; Stevenson-Holt et al. 2014). Each landscape component is a numeric variable, difficult to express in terms of real resistance values, mainly due to lack of knowledge of dispersion forms, and consequently, in the gene flow.

To evaluate the potential resistance that the landscape imposes on gene flow among Baru populations, and to reveal information on the values of the parameters and the processes behind the pattern of genetic divergence between populations. We used Pattern-Oriented Modeling (Grimm et al. 1996; Grimm et al. 2005; Grimm and Railsback 2012; Topping et al. 2012), an approach based on the genetic divergence between “Baru” populations. Pattern-Oriented Modeling (POM) allows finding which combination of parameters maximize the correlation between genetic and dispersal, and subsequently, the selection of the “best” resistance surface under this relationship.

To build the resistance surfaces, resistance values in the interval between 0 and 100, were randomly assigned to each landscape components, generating 10.000 surfaces with different values of resistance for each class. Despite the randomization of resistance values, the landscape configuration (classes) has not changed, remaining constant in all generated surfaces (Figure 2).

**Fig. 1.**
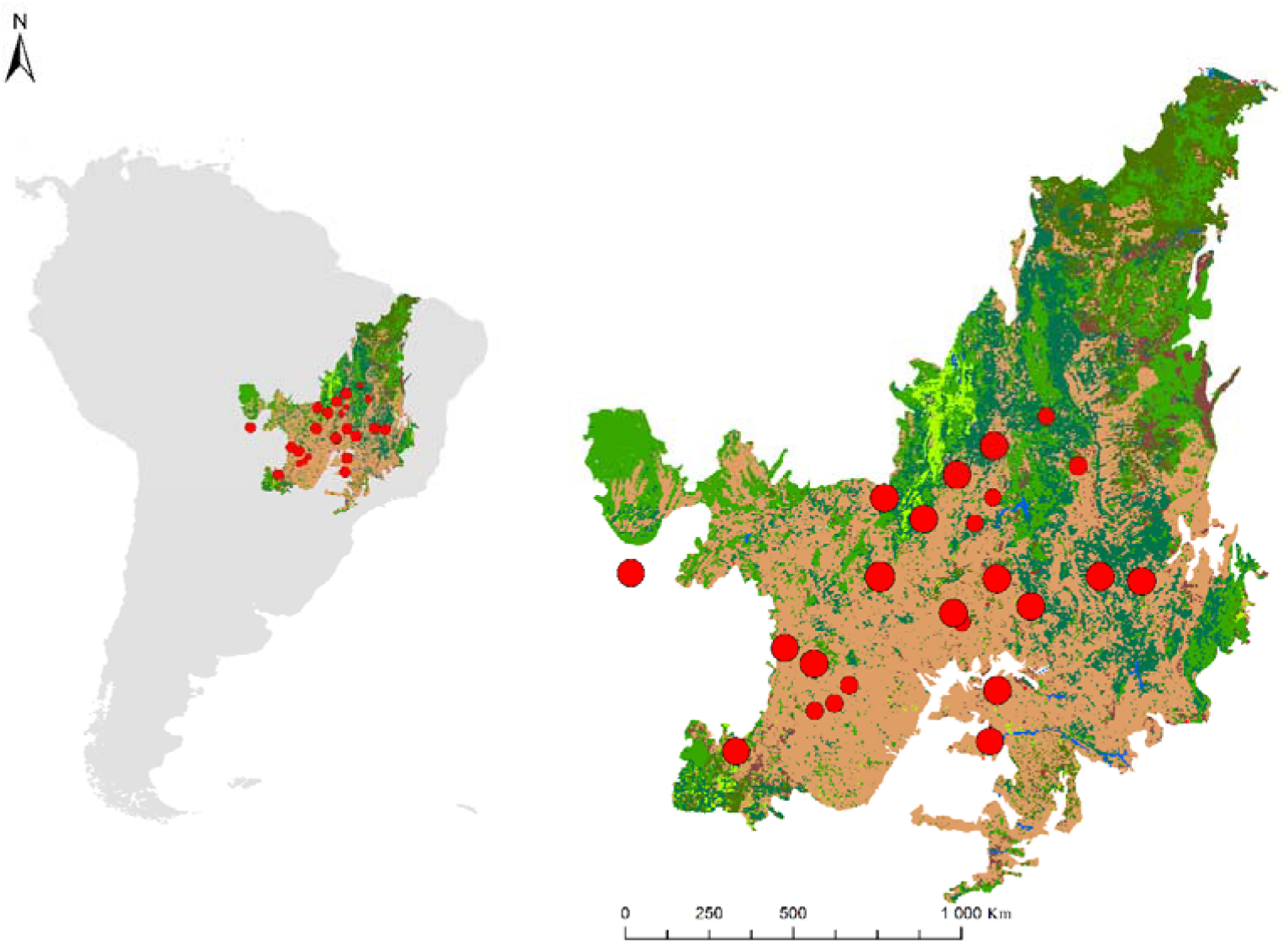
Distribution of *Dipteryx alata* population studied in the Brazilian Cerrado, with the landscape components (classes) analyzed in resistance surface. Areas in green tones are natural areas, and brown tones are predominantly disturbed areas. Red circles indicate the number of 25 populations genotyped, ranging between 12 and 37 individuals

**Fig. 2.**
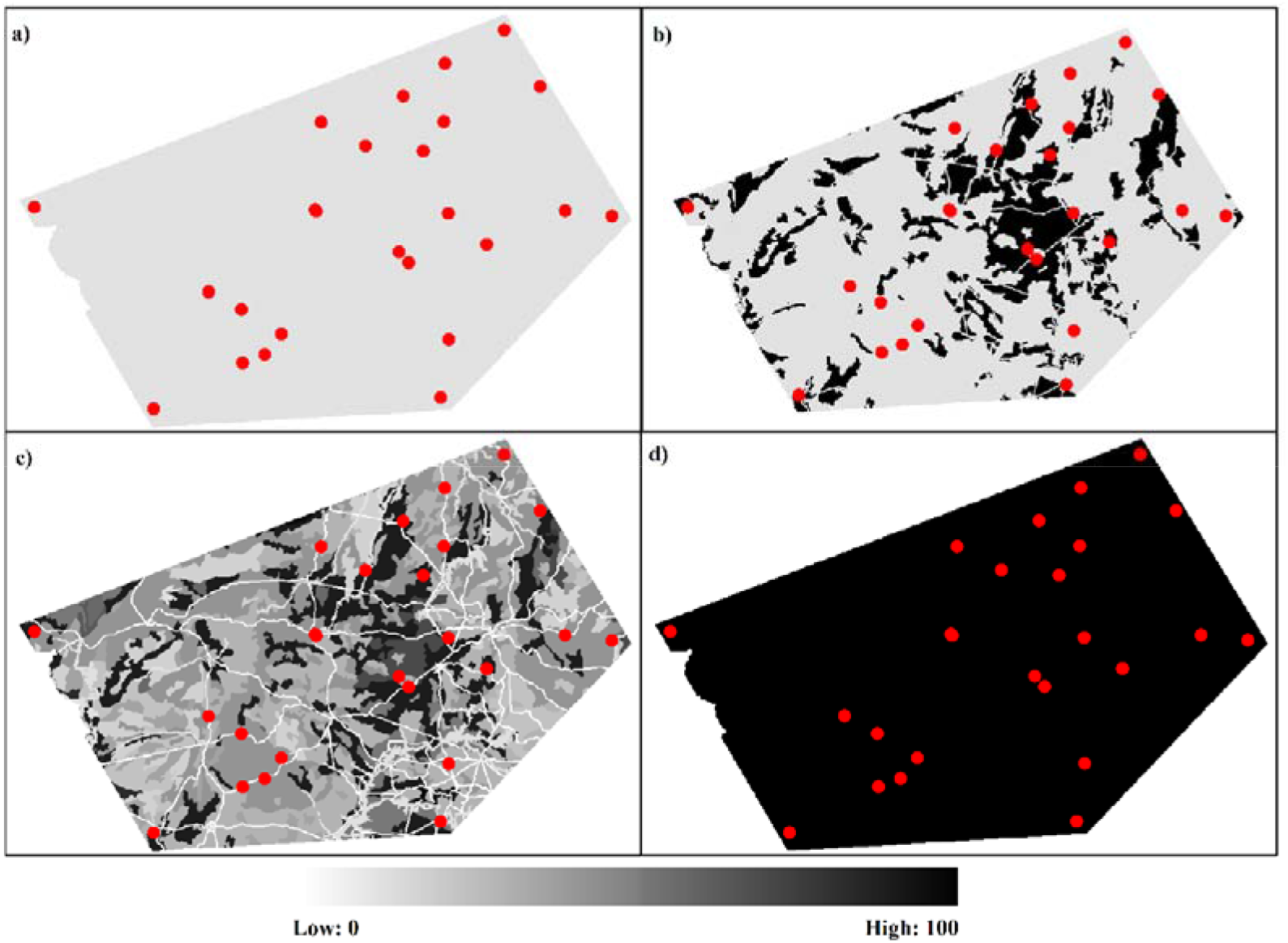
**a)** and **b)** are homogeneous surfaces with respectively resistance values of 0 and 100 in all areas. The surfaces **c)** and **d)** are heterogeneous and contain the parameterized classes with values of 0 to 100 at intervals of 1

We started the modeling with a “null” model, where all cells on the resistance surface were zero, equivalent to considering only the Euclidean distance between pairwise populations. Next, we created more complex landscape surfaces incorporating heterogeneity, with randomized values of resistance in each of its components and comparing them with the null model. In a third cycle, we systematically removed the complexity of the surfaces in the selected models. We evaluated the importance of each class and its resistance in the model and validation. Our work emphasizes the second step of the POM, the estimation of parameters in the creation of resistance surfaces. Our most significant interest is to quantify the resistance imposed by each landscape component to the dispersion and, consequently, to the genetic diversity and gene flow between populations.

We generate 10.000 resistant surfaces and calculate the effective resistance matrices for pairs of populations according to the Electrical Circuit Theory (McRae 2006; McRae and Beier 2008 McRae et al. 2008). Differently from the more commonly used least-cost path, which indicates a single possible connection path among populations, the effective resistance calculation is based on the Electrical Circuit Theory, allowing calculating the resistance distances considering multiple pathways (McRae et al. 2008). The resistance calculation based on the Electrical Circuit Theory was generated through Circuitscape software (v.4.0 Beta-; McRae, 2006). We used the ResistanceGA package in R (Peterman 2014) to carry out the analysis in Circuitscape.

### Spatial Analysis

We correlated the spatial arrangements (in terms of resistance distance between populations) with the F_ST_ pairwise using different Mantel tests (see Diniz-Filho et al. 2013, for a recent review). The significance of Pearson matrix correlations was estimated from 999 permutations.

To check the influence of landscape resistance on the species’ gene flow, maintaining the effect of geographic distance between populations permanent, we also used a partial Mantel test (Legendre and Legendre 2012), calculated using R software (R Core Team 2014) with *mantel.partial* function of the vegan package (Oksanen et al. 2012). The Akaike information criterion (AIC) was used to select the best (most satisfactory) models (deLeeuw 1992), which were considered those with ΔAIC values below three. The ΔAIC values were also used to calculate the AIC weight in each model (Akaike weight - Wi) (Burnham and Anderson 2002; Diniz-Filho et al., 2008), revealing the level of certainty in achieving the best explanatory model against all the other tested models.

Each landscape component has an individual contribution to the explanatory power of models. To assess this individual contribution of each landscape component in all select models (equation 1),

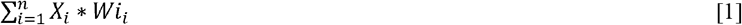

we multiplied (wi) by the resistance values of each landscape component (Xi) in the selected models (n). With this approach, we were able to measure the influence of each landscape component on gene flow, translated in genetic distances observed.

We then used the mean resistance of selected models to create a final current map to verify possible species dispersion routes and identify the areas that most contribute to connectivity between populations (as seen in Castilho et al., 2011; McRae, 2006; McRae et al., 2008). The value of each cell in the landscape represents an amount of current flowing through it, analogous to the percolation of the species (McRae et al. 2008). Where the landscape is less resistant, the routes of dispersion are more likely to occur.

## Results

We found a significant amount of genetic divergence between populations, with an average F_ST_ of 0.258 to establish a comparison among all the populations. The genetic divergence based on F_ST_ values was positively correlated with the geographic distance (Mantel test, r = 0.4805; P <0.001), showing a linear relationship between genetic and geographic distances in which genetic divergence between populations (inverse of F_ST_) increase with geographic distance increase (Figure 3).

**Fig. 3.**
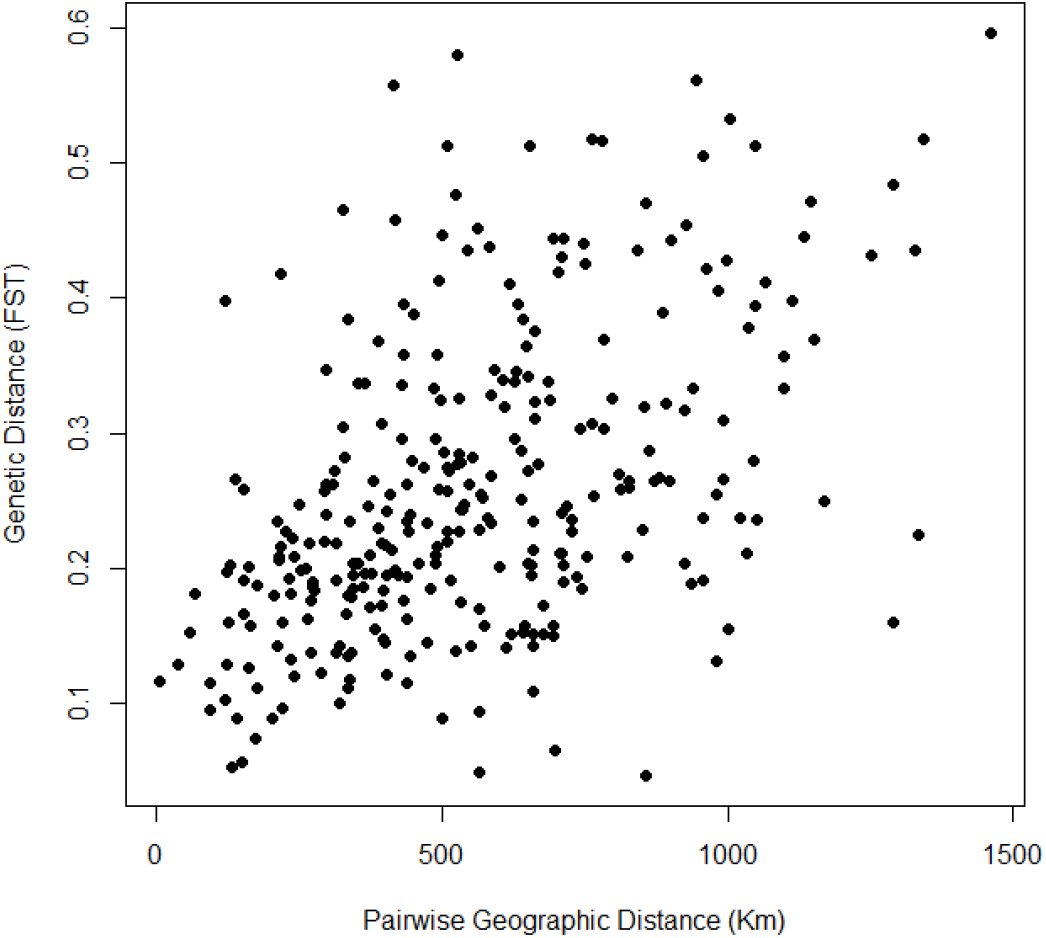
Relationship between pairwise FST and geographical distance

For the resistance surfaces generated, Mantel correlation coefficients ranged between −0.2019 and 0.6736 (Figure 4). About 56% out of the 10.000 resistance surfaces have a higher correlation with F_ST_ than the one obtained with geographic distance alone. Partial Mantel was used to taking into account the effect of geographical distance, providing thus an estimate of how genetic divergence is explained by landscape alone. The average partial Mantel correlation coefficient was equal to 0.2016, demonstrating explanatory gain for genetic divergence among populations, and increase their respective resistances in the model.

**Fig. 4.**
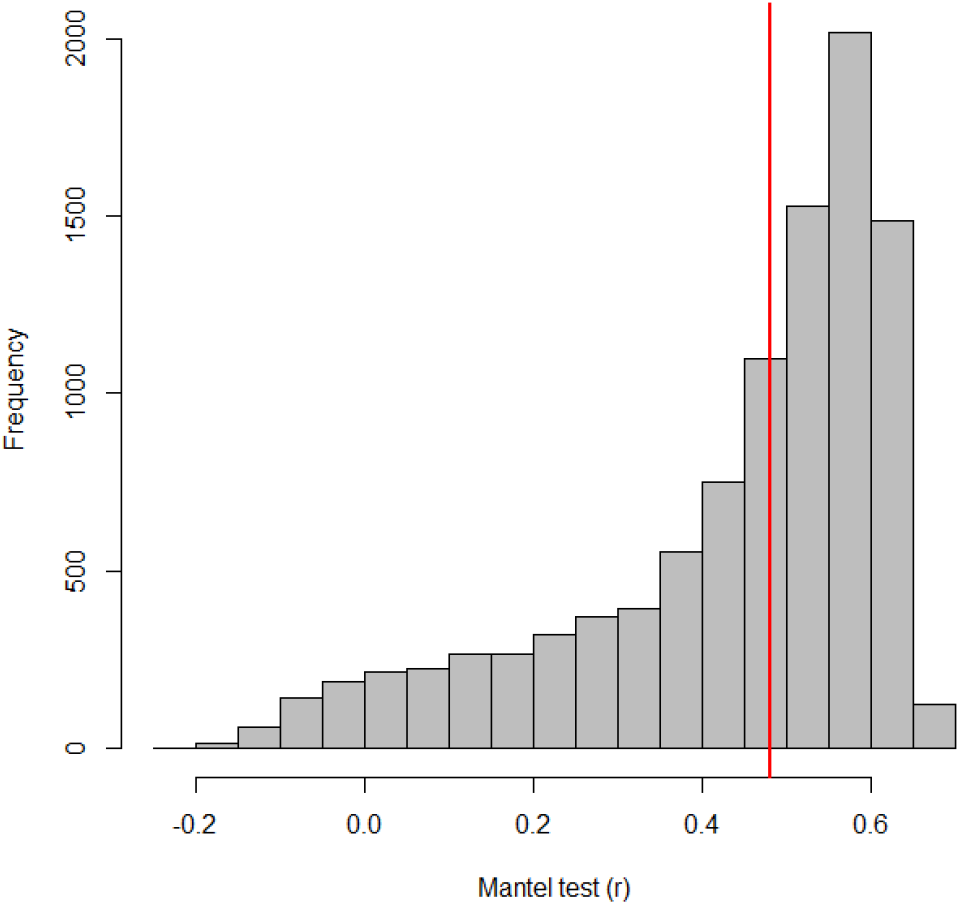
Histogram of Pearson’s correlation coefficient (r) values obtained using the Mantel test between pairwise FST and resistance matrices, highlighting the correlation coefficient between the FST and geographical distances, r = 0.4805 (red line)

Three out of the 10.000 resistance surfaces were selected by ΔAIC under 3. These models are providing the best relationship of landscape with the genetic diversity among the “Baru” populations. The Mantel between landscape resistance and genetic divergence suggest a major adjustment of resistance models, about r = 0.67, much larger than a pure geographical model, with r = 0.48 (Table 1). All Mantel tests were significant at P <0.001, with 999 permutations. The relative Wi suggests that there is about a 30% chance that the surface S-9247 is the more adjusted, a high chance since 10.000 models were generated. The second and third models selected have a reduced chance of 8% and 6.8%, respectively.

**Table 1.** Resistance surface selected according to ΔAIC.

| Surface | Mantel | Partial Mantel | RT | RR | RG | a | b | c | ΔAIC | Wi |
| --- | --- | --- | --- | --- | --- | --- | --- | --- | --- | --- |
| S-9247 | 0.6736 | 0.5404 | 0.456 | 0.454 | 0.23 | 0.225 | 0.228 | 0.001 | 0 | 0.29 |
| S-3939 | 0.6702 | 0.5364 | 0.452 | 0.449 | 0.23 | 0.221 | 0.227 | 0.002 | 2.528 | 0.082 |
| S-4515 | 0.6697 | 0.5319 | 0.449 | 0.449 | 0.23 | 0.218 | 0.23 | -0.001 | 2.895 | 0.068 |

In partial mantel, the resistance distance explains approximately 45% of the genetic divergence between populations - RR, more than twice the geographical distance explanation provided - RG (23%) (Table 2). By partitioning the effects, about 49.2% out of all the genetic divergence is explained only by resistance (a), about 50.7% of the strength by the overlap with geographical distance (b), and 0.1 % is defined exclusively by the geographic distance (c).

**Table 2.**
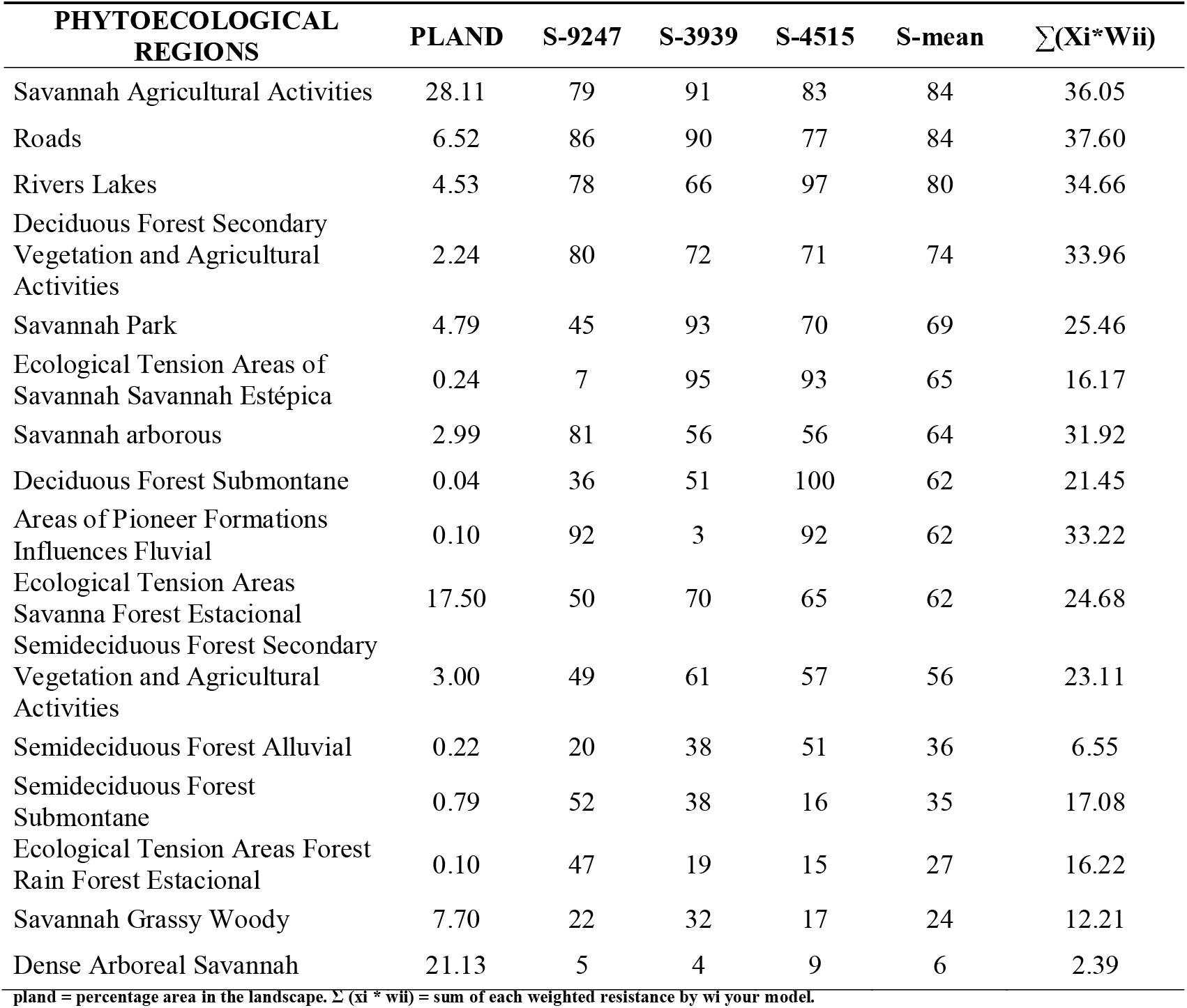
Influence of landscape features on the gene of flow *Dipteryx alata.* Costs on the resistance surfaces selected and the sum of the pondered cost per wi.

The study area is composed of 16 different classes (Table 2). With 28% of the landscape composed by Savannah Agricultural Activities, 21% of Dense Arboreal Savannah, 17.8% of a transition area between different vegetation types (Ecological Tension Areas), 7.7% of Savannah Grassy Woody, 6.5% of roads and 4.5% of rivers and lakes. Other classes of landscape account for the remaining 14.5%. The models selected suggest that areas with lower resistance are those with vegetation types Dense Arboreal Savannah and Savanna Grassy Woody (grassland). Savannah Agricultural Activities, Roads, and rivers cause higher resistance.

We built a final effective resistance surface from the weighted resistance costs and the weight of the selected models (Figure 5a). The surface is correlated with the genetic divergence of “Baru” populations, with r = 0.6854 (P <0.001). The primary gene flow routes of the “Baru” occur where the landscape is less resistant, and dispersal routes are more likely to occur (Figure 5b). The models suggest that areas with lower resistance to gene flow are the Savanna and Savanna Dense Arboreal Grassy Woody. Roads, rivers, and the agricultural regions of Cerrado cause higher resistance to gene flow. We used a current map to preview important connectivity areas between populations; in this case, the minor resistance areas represent areas propitious for species life and to areas suitable to the disperser animals of the species. Warmer colors (purple and red) indicate areas with less current density; areas with higher connectivity are shown in yellow.

**Fig. 5.**
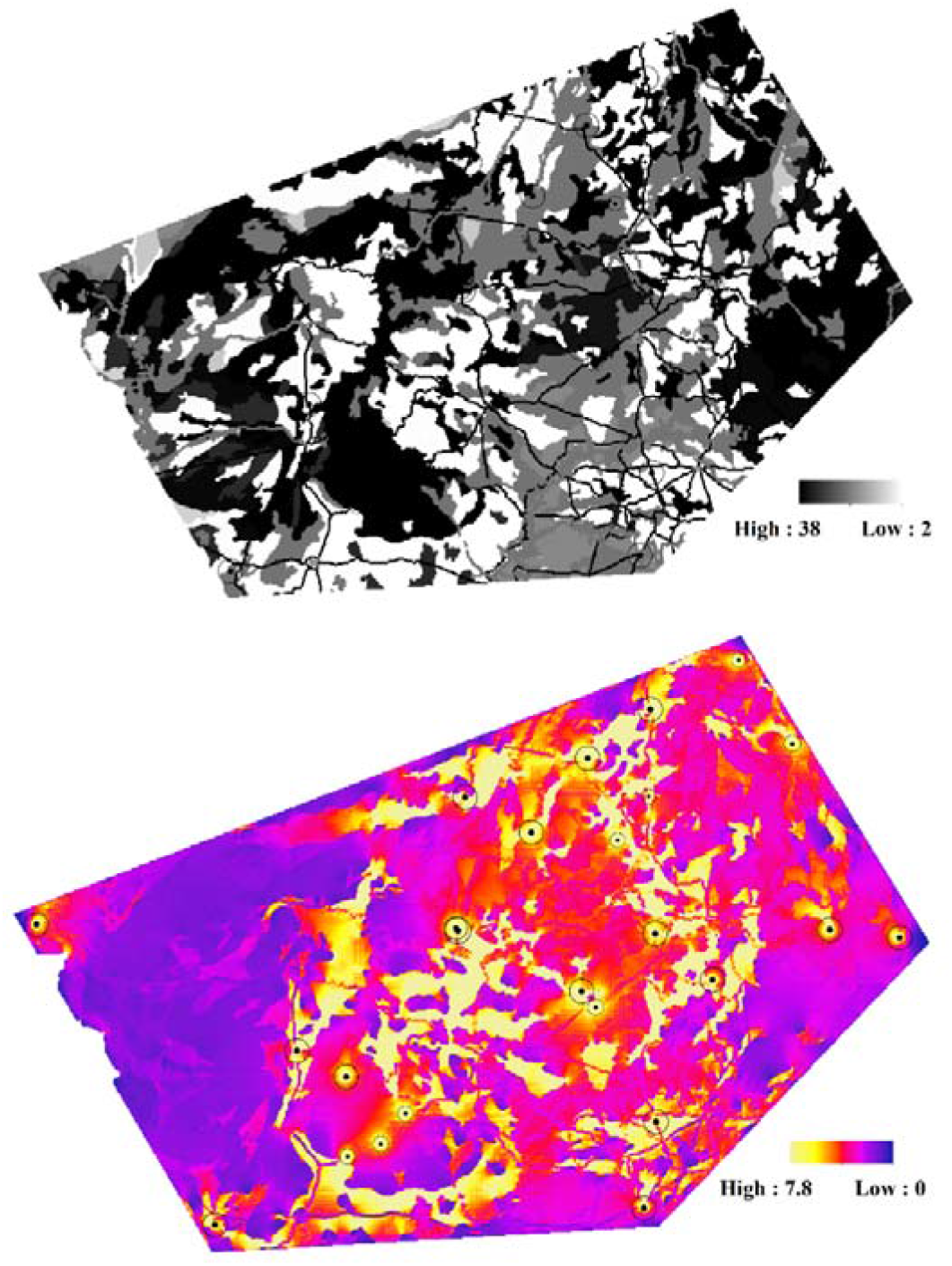
**a)** landscape resistance map for Baru (*Dipteryx alata*) in the Brazilian Cerrado with average resistance costs ranging between 2.3928 and 37.587. The resistance gene flow was parameterized with average model parameter estimates for these variables. The lighter areas have lower resistance, and the darker has a higher resistance. **b)** current map with the main routes of lower resistance to the dispersion of Baru

## Discussion

In population genetics, several approaches have been used to investigate patterns and infer microevolutionary processes involved in population differentiation. Landscape genetics is the study of how landscape pattern (the distribution of suitable habitat, barriers, etc.) affects gene flow and genetic differentiation of species (Holderegger and Wagner 2008; Manel and Holderegger 2013). Here we correlated F_ST_ with geographical distances and landscape features using distinct approaches that allow us to decouple their effects. Our results seem to be sufficiently robust to can furnish a description of landscape influence in the genetic variability in a relatively well-known species and corroborate previous analyses of population genetic structure with “Baru” populations (e.g., Soares et al. 2008; Telles et al. 2014), adding and complementing information, suggesting influences of anthropic actions among these populations. The Circuitscape has been widely used to check used how landscape features influence genetic connectivity (Adams et al. 2016, Mateo-Sánches et al. 2015 and Pérez-Espona et al. 2012). For creating the resistance surface is necessary parameterizing cost surfaces by assigning weights to different landscape elements has been challenging, however, because real costs are rarely known (Koen et al. 2012). Here, we seek to understand how each landscape components influence genetic divergence between populations of *Dipteryx alata.* Using a Pattern-Oriented Modeling approach, we established which cost configuration best explains the genetic divergence between the studied populations, bringing The approach we used allows getting more insights into the importance of landscape to the genetic flow of populations based on a new way to minimize the arbitrariness in the parameterization of resistance surfaces.

The patterns found in this study show the benefits of using an additional set of information to create these surfaces and to interpret the genetic differentiation among populations. Even though another study will have been trying to develop ways to measure the resistance surfaces more clearly by using modeling (e.g., Shirk et al. 2010; Spear et al., 2010) until now, there is no consensus on the most effective approach. Spear et al. (2005) took a big step by using the lowest cost path analysis and discussing the difficulty of developing cost parameters for different habitat types, without having the necessary data (the species biology information and its dispersion). But they did not use cover type because they did not use data that would allow quantifying the specific numerical cost of moving through of the studied salamander species. Koen et al. (2012) carried out a sensitivity analysis of three methods to parameterize a cost surface and two models of landscape permeability. They check that developing a cost surface improves the accuracy of functional connectivity estimates, especially when cost weights are selected through statistical model fitting procedures.

The “Baru” has been widely studied in ecological and genetic terms due to its economic, environmental, and cultural importance. Our results seem to be sufficiently robust to can furnish a description of landscape influence in the genetic variability in a relatively well-known species. Our results corroborate and expand our understanding of the factors driving the population genetic structure of this species. Soares et al. (2008) detected that in local populations situated at short geographic distances has been the spatial structure of genetic divergence demonstrated a pattern of genetic discontinuities, suggesting influences of anthropic actions among these populations. Other studies with this species (e.g., Telles et al. 2014; Diniz-Filho et al. 2015; Soares et al. 2015) have begun to worry about the influence of environmental characteristics on the genetic differentiation of “Baru” populations. Telles et al. (2014) correlate different landscape metrics with the genetic divergence of these same Baru populations finding a strong correlation between the percentage of natural remnant and genetic divergence of populations, demonstrating how human occupation had effects such as habitat loss and fragmentation. Soares et al. (2015) and Smith et al. (2015) discuss and analyze the distribution of these populations from a *center*-periphery dynamic, using this environmental suitability and demonstrating a historical influence on the distribution of these populations.

The genetic divergence between populations of D. *alata* is better explained by landscape structure than by merely geographic distance. Only about 23% of the genetic divergence is explained by geographic distance, reinforcing that factors other than the geographical distance influence the genetic differentiation among Baru populations. For the resistance surfaces generated, Mantel tests resulted in correlation coefficients varying between −0.2019 and 0.6736, demonstrating the importance of considering landscape components. While the Euclidean distance was a path for this system, this is not usually the best case for studies that have landscape additive information (Coulon et al. 2004, Emel, and Storfer 2015).

Regarding landscape components, forest formations did not have a significant influence on the results, mainly due to its small presence in the landscape. The landscape classes that occur in small quantity facility the transposable for “Baru” dispersers, such as birds (macaws) and mammals (monkey, agouti, and livestock), all long-distance dispersers (Ribeiro et al. 2000). Savanna arboreal dense and savanna grassy woody (grassland) behaved as classes with lower resistance to species, observing that low resistance has a direct association with high percolation for animals that disperse their seeds and pollen, as it is a plant with zoochoric dispersion. We expected low resistance in savanna arboreal dense and savanna grassy woody since this species can be found in this type of environment. The classes that showed higher resistance to “Baru” dispersion were the savanna agricultural area, rivers, lakes, and roads, all of them with values close to 35, suggesting high resistance, but also high percolation since it can vary between 0 and 100 (35 is medium resistance). We expected that these landscape classes to have higher resistance considering that they are disturbed areas. We believe that agricultural areas cause increased resistance, mainly due to its instability plant; the Savanna case, they constitute a very high percentage of areas, 28.11%, of the entire study area, with long stretches. The fact that dispersion is mostly associated with terrestrial animals and large and medium-sized flying animals justify the median interference of landscape on species’ gene flow. We expected that species with more restrictive dispersal to have more extreme and high values, which reflects mainly on the ability of “Baru” dispersers to overcome barriers and high resistance areas along the dispersion process. The relative importance of landscape components and their spatial patterns can be the key for identifying their influence in microevolutionary processes driving population divergence.

From a conservation point of view, the current map (McRae et al. 2008) is a source of information on population connectivity. It has been used to demonstrate critical areas for species’ connectivity maintenance (Castilho et al. 2011 and Schwartz et al. 2009). This enhances the understanding of isolated species and facilitates the process of decision-making regarding the main routes of connectivity between populations. Therefore, this work shows an improvement regarding the previous analyses by demonstrating the influence that landscape components have on the processes that generate such genetic variation. Further studies comparing different tree species in this region would allow correlating the weights obtained with their life-history attributes, reinforcing the interpretation of how these differences are captured by IBR and IBE models. Moreover, once these relationships are better established, it would be possible to evaluate how the profound ongoing landscape changes in Brazilian Cerrado (Bonanomi et al. 2019) would disrupt gene flow and, consequently, would lead some economically important species as the “Baru” to local or global extinction.

## Declarations

### Availability of data and material

The data used in work will be made available after approval of the article by Journal

### Code availability

The custom code created and used in the development of the work will be made available after the Journal approved the manuscript.

### Authors’ contributions

All authors contributed to the study conception and design. Material preparation, data collection were performed by Kelly Souza; Mariana Pires de Campos Telles; Thannya Nascimento Soares; Lazaro José Chaves. Analysis was performed by Kelly Souza; Jesús N. Pinto-Ledezma; Clarissa Bonafé Gaspar Ruas; Ricardo Dobrovolski; José Alexandre Felizola Diniz-Filho. Kelly Souza wrote the first draft of the manuscript, and all authors commented on previous versions of the manuscript. All authors read and approved the final manuscript.

## Electronic Supplementary Material

**Table S1:** Geographical coordinates of Baru Populations

| Population | City / State | Longitude | Latitude | Number of Samples |
| --- | --- | --- | --- | --- |
| 1 | Cocalinho/MT | -14.397 | -50.996 | 32 |
| 2 | Nova Nazaré/MT | -13.839 | -52.042 | 32 |
| 3 | Pirenópolis/GO | -15.997 | -49.034 | 32 |
| 4 | Sonora/MS | -17.853 | -54.704 | 31 |
| 5 | Alcinópolis/MS | -18.268 | -53.926 | 32 |
| 6 | Alvorada/TO | -12.449 | -49.115 | 32 |
| 7 | São Miguel do Araguaia/GO | -13.225 | -50.103 | 32 |
| 8 | Luziânia/GO | -16.732 | -48.13 | 32 |
| 9 | Icém/SP | -20.347 | -49.216 | 31 |
| 10 | Monte Alegre de Minas/MG | -18.978 | -49.024 | 32 |
| 11 | Estrela do Norte/GO | -13.828 | -49.14 | 12 |
| 12 | Santa Terezinha de Goiás/GO | -14.52 | -49.628 | 12 |
| 13 | Arinos/MG | -15.934 | -46.273 | 32 |
| 14 | Pintópolis/MG | -16.061 | -45.166 | 32 |
| 15 | Chapadão do Sul/MS | -18.846 | -52.982 | 13 |
| 16 | Água Clara/MS | -19.33 | -53.377 | 13 |
| 17 | Camapuã/MS | -19.528 | -53.9 | 13 |
| 18 | Indiara/GO | -17.162 | -49.973 | 13 |
| 19 | Aragarças/GO | -15.9112 | -52.187 | 27 |
| 20 | Aragarças/GO | -15.9482 | -52.158 | 37 |
| 21 | Palminópolis/GO | -15.9121 | -50.201 | 32 |
| 22 | Chapada da Natividade/TO | -11.6614 | -47.714 | 12 |
| 23 | Arraias/TO | -12.9904 | -46.863 | 15 |
| 24 | Anastácio/MS | -20.611 | -56.004 | 31 |
| 25 | Porto Esperidião/MT | -15.853 | -56.8222 | 30 |

